# Genetic variation in behavioral and physiological responses to copper in *Drosophila melanogaster*

**DOI:** 10.64898/2026.08.23.746539

**Authors:** Md Meftahul Zannat, Jordan C. Jones, Maggie Ridgway, Elizabeth R. Everman

## Abstract

Anthropogenic copper (Cu) contamination from agriculture, mining, and industrial runoff creates environmental gradients affecting physiology and behavior in wild populations. While Cu toxicity in *Drosophila melanogaster* is well characterized, it remains unclear whether Cu resistance is one integrated trait or several independently evolving components. Using a subset of recombinant inbred lines (RILs) from the *Drosophila* Synthetic Population Resource (DSPR), we measured three components of Cu response: feeding avoidance, oviposition avoidance, and physiological tolerance (median lethal time, LT_50_) under sustained Cu exposure. All three traits showed substantial phenotypic variation among RILs. Feeding and oviposition avoidance were both highly heritable (H² ∼ 0.88), and RIL identity accounted for 49.5% of the variance in LT_50_. However, the three traits showed no significant correlation across RILs, indicating distinct genetic architecture. We identified a single male specific quantitative trait locus (QTL) on chromosome 2R that explained 17.7% of the variation in feeding preference; the interval included candidate detoxification genes *Jheh1*, *Jheh2, Jheh3* and *sano*, the latter of which is associated with olfactory behavior. No significant QTL were detected for oviposition preference, suggesting a highly polygenic structure that may difficult to detect with our limited panel size. Together, these results indicate that Cu resistance in *D. melanogaster* is genetically modular. Behavioral avoidance during feeding, oviposition, and physiological tolerance are heritable but architecturally distinct components, each with potential to respond to selection independently.

## Introduction

Stressors may have immediate effects that influence an organism’s health and/or long-term effects that alter the evolutionary trajectory of populations. Heavy metals are one such class of stressors, persisting in terrestrial and aquatic ecosystems after being released from mining, industrial discharge, and agricultural effluents (Scott and Sloman 2004; Ali et al. 2019; Abd Elnabi et al. 2023; Honorio et al. 2023; Mititelu et al. 2025). Metals like copper (Cu), lead (Pb), and cadmium (Cd) may accumulate in soil and food sources where they affect organisms across several biological levels. On a molecular basis, higher concentrations disrupt redox balance, generate reactive oxygen species (ROS), and damage proteins and nucleic acids (Stohs and Bagchi 1995; Nofal et al. 2023; Rasmy et al. 2024). These biochemical effects result in reduced growth, fecundity, and survival, ultimately shaping the evolution of exposed populations (Gordon 2003).

However, environmental abundance of toxic elements doesn’t determine exposure independently. Feeding behaviors, habitat choice, or oviposition site selection may influence an individual’s exposure to toxic substances (Smith et al. 2007; Peterson et al. 2017; Burden et al. 2019; Araújo et al. 2020; Jacquin et al. 2020). For example, aquatic invertebrates such as *Daphnia magna* reduce feeding rates and avoid contaminated water when exposed to heavy metals, resulting in lower contaminant intake (Lampert, W. 1987; Barata et al. 2002). Similarly, freshwater fishes alter their habitat use and feeding behavior to avoid polluted environments, demonstrating that behavioral responses can reduce toxic exposure (Jacquin et al. 2020; Brand et al. 2025; Shahriar et al. 2025). Behavioral avoidance can therefore be regarded as one component of toxicological resistance rather than as an independent process.

Notably, individuals that appear physiologically resistant may simply ingest less toxin (Sparks et al. 1989; Scott and Sloman 2004; Zalucki and Furlong 2017). A recent study in *Drosophila melanogaster* illustrates this coupling directly; insecticide resistance mediated by *Cyp6g1* is associated with avoidance of DDT-containing oviposition substrates, suggesting that physiological resistance and behavioral avoidance can evolve together (Alves et al. 2026). Whether this coupling is a general feature of toxin resistance or specific to xenobiotics and their detoxification pathways remains unclear. This question is especially important for heavy metals such as Cu, which are harmful at high concentrations but essential at low levels.

For more than a century, the common fruit fly, *Drosophila melanogaster*, has been a key component of genetic research, providing insight into gene function (Morgan 1910; Bellen et al. 2010; Mackay et al. 2012), environmental stress response (Rand 2010), and genetic variation in physiology and behavior (King et al. 2012). Recently, *D. melanogaster* has been utilized to investigate the effects of chemical exposures on learning, memory, and other behaviors (Welch and Mulligan 2022). Metals like iron (Fe), zinc (Zn), and Cu are micronutrients of the fly diet; however, they can become toxic when intracellular concentrations exceed homeostatic capacity (Balamurugan et al. 2007; Calap-Quintana et al. 2017; Navarro and Schneuwly 2017; Abolaji et al. 2020). At higher concentrations, such imbalances can disrupt critical cellular processes, neural signaling, and behaviors (Bahadorani and Hilliker 2009; Bonilla-Ramirez et al. 2011; Monnier et al. 2018; Tran Thanh et al. 2024).

Among these metals, Cu provides an excellent system for studying how organisms balance an essential element with toxicity. Cu is required as a cofactor for enzymes such as cytochrome c oxidase and Cu/Zn superoxide dismutase, but at higher concentrations it generates ROS and impairs neural function (Balamurugan et al. 2007; Bahadorani and Hilliker 2009; Bonilla-Ramirez et al. 2011; Abolaji et al. 2020). Cu pollution can enter terrestrial and aquatic systems through fungicide use, industrial smelting waste, and urban runoff, creating local hotspots that reduce invertebrate survival and reproduction (Michaud and Grant 2003; Wightwick et al. 2010; Zubrod et al. 2014; Husak 2015; Ge et al. 2023). In *D. melanogaster*, Cu exposure reduces feeding, fecundity, and locomotor activity, and gravid females generally avoid laying eggs on Cu supplemented substrates (Bahadorani and Hilliker 2009; Scott 2018; Budiyanti et al. 2022). However, it remains unclear to what degree Cu avoidance varies among genotypes, whether this variation is heritable and genetically linked with physiological tolerance.

Together, feeding and oviposition are complex behaviors influenced by genetic variation and by chemosensory, gustatory, and neural processing (Mackay et al. 2012; Scott 2018). The gustatory receptor gene family in *Drosophila* consists of 68 genes, many of which are involved in detecting bitter or aversive compounds (Scott 2018). Oviposition site choice also involves mechanosensory cues and neuropeptide signaling, including *ILP7* expressing neurons that regulate egg-laying (Yang et al. 2008). These pathways are relevant to Cu responses and may shape how Cu is detected, avoided, and tolerated. Lee et al., (2010) found that, *Drosophila* gustatory receptor neurons (*Gr32a*, *Gr33a*, and *Gr66a*) can detect aversive compounds, whereas ionotropic receptors mediate the detection of salts and divalent cations, including calcium (Ca^2+^) (Puri and Lee 2021). The role of these receptors in Cu detection is unknown, and the genetic basis of metal avoidance remains poorly described.

To address this gap, we used a subset of strains from the *Drosophila* Synthetic Population Resource (DSPR) (King et al. 2012) to characterize genetic variation in Cu avoidance and tolerance and to identify genomic regions associated with that variation. We focused on three complementary phenotypes, each representing a different component of resistance. Feeding avoidance and oviposition avoidance reflect distinct behavioral decisions. Feeding is shaped largely by gustatory perception and digestive signals, whereas oviposition site choice reflects a maternal decision guided by chemosensory and mechanosensory cues (Yang et al. 2008; Bahadorani and Hilliker 2009; Gou et al. 2014). Physiological tolerance under continuous Cu exposure, measured as median lethal time (LT_50_), reflects the ability to endure Cu independently of behavioral avoidance. Together, these traits allow us to determine whether Cu resistance is caused primarily by avoiding the toxin, tolerating it, or both. If behavioral avoidance and physiological tolerance share genetic pathways, we expect a positive correlation and shared quantitative trait loci (QTL). Alternatively, if they are controlled by different biological pathways, the traits may be heritable but weakly correlated, with largely distinct genetic architectures. This distinction is significant beyond *Drosophila*. If resistance is modular, selection may act on its components independently, allowing populations to evolve different responses as heavy metal pollution varies geographically and temporally.

## Methods

### Fly RILs and rearing

We focused on a subset of recombinant inbred lines (RILs) from the *Drosophila* Synthetic Population Resource (DSPR). Feeding response was measured in 197 DSPR RILs. Oviposition and physiological estimates of Cu response were measured in 155 and 20 DSPR RILs, respectively. Prior to testing, all flies were reared under standard conditions (25°C, 12:12 light-dark cycle, relative humidity ∼50%) on cornmeal-molasses-yeast diet to ensure their health and prevent any heavy metal exposure prior to the assay. Flies were maintained in rearing conditions for all experiments described below.

### Microplate Feeding Assay

Polystyrene 96 well plates were prepared with 1.5% agarose starvation medium (*Drosophila* agar, water, propionic and phosphoric acid preservatives). While the medium was in a liquid state, 100uL were added to each of the 96 wells with a multichannel pipet. Flies (3-5 days old) were transferred under CO_2_ anesthesia to the starvation media plate (1 fly/well) attached to a 3D printed coupler which allowed each fly to gain access to food after fasting (Walters et al. 2021). Two Rows of the plate were left empty to correct for evaporation of liquid food. We assessed one RIL per 96 well plate and up to 36 flies per sex. Once sorted, the flies fasted for 24 hours.

Liquid food was prepared on the day of the experiment by combining 2.3g of sucrose with 1.1g Bacto Yeast Extract (Thermofisher scientific C212720), 52mL dH_2_O, and 224uL of 10 mg/mL blue dye stock (erioglaucine disodium salt; EDS, Sigma 861146) in a master mix. Control food was prepared by adding an additional 1mL of dH_2_O to 13mL of the master mix to bring the total volume to 14mL. For the 2mM Cu food preparation, 13mL of the master mix was combined with 440uL dH_2_O, and 560uL 50mM CuSO_4_ (total volume = 14mL) (Cu (II) Sulfate; Sigma-Aldrich C1297). Once both control and experimental food were prepared, 10uL of each food type were dispensed into the 1536 well plates utilizing a multichannel pipet as arrayed in (Walters et al. 2021). A film (Seal Plate Sealing film, Genesee #Cat 12-167) was placed over the top and bottom, sealing the 1536 well plate, and plates were briefly centrifuged. Absorbance values (630nm) were taken for the 1536 well plate before the flies were allowed to feed. To allow the flies access to the food, the film was perforated in each well containing the food options. After the flies had access to the food for 24 hours they were removed, and a second absorbance reading was taken of each 1536 well plate. The 96 well plates were inspected and any flies who did not survive were excluded from data analysis.

Cu feeding preference was determined by measuring the difference between the initial and final absorbance readings of the 1536 well plate containing both control and experimental Cu supplemented food after accounting for evaporation (Walters et al. 2021). As food was consumed by flies, absorbance values decreased providing an estimate of consumption. Negative feeding preference values indicated avoidance of Cu food; positive values indicated a preference for Cu food.

Visual inspection of residuals from the ANOVA model (Preference_2mM ∼ patRIL × Sex) indicated a right skewed distribution. Because the sample was large (n = 10,479) and exceeded the upper limit recommended for the Shapiro-Wilk test (n ≤ 5,000), we did not apply this formal test of normality. Levene’s test indicated heteroscedasticity F _(378, 10101)_ = 3.0, P < 0.00001), although the magnitude of variance heterogeneity was modest and likely amplified by the large sample size. To improve residuals normality and reduce the influence of right skewness on parametric inference, feeding preference was square root transformed as (sqrt (Preference 2mM + 1)); a constant of 1 was added prior to transformation to accommodate negative preference values. We then refit the ANOVA on the transformed response variable and report results from this model. Sex-specific heritability was estimated using a linear mixed effects model and the *varcomp* function from the *nlme* (Pinheiro et al. 2025) and *ape* (Paradis and Schliep 2019) packages in R (R Core Team 2025). We performed QTL mapping of Cu aversion for males and females using the DSPRqtl version 2.0-5 and DSPRqtlDataB version 2.0-1 respectively (King et al. 2012). All QTL analyses were performed in R (R Core Team 2025).

### Oviposition assay

We conducted the oviposition assay using custom built acrylic oviposition chambers fabricated by the University of Oklahoma Innovation Hub. The chamber design was based on Gou et al. (2016), with modifications to well geometry and chamber sealing that improved fly retention and reduced cross contamination between substrate types (Supplementary Figures S1, S2). Each chamber consisted of a single transparent acrylic plate containing two identical arrays of 30 individual wells (5 rows × 6 columns per array). Wells were rectangular and separated by thin acrylic dividers to prevent cross contamination of eggs or substrates. Two additional transparent acrylic plates were placed above the well plate to provide unobstructed access for flies, allowing them to freely choose between control or Cu substrates and lay eggs in either well. The plates were aligned and secured using six threaded bolts.

We prepared two types of egg-laying media using the semidefined medium recipe (Backhaus et al. 1984). Control media was prepared according to the standard recipe. We supplemented the media with 10 mM CuSO₄, a concentration that maintained >90% adult survival while eliciting robust avoidance in pilot oviposition assays (mean preference index = −0.21). We allowed substrates to solidify and used them one day after preparation.

To standardize mating status, females were kept with males for the oviposition experiment. For additional protein and to encourage oviposition, we also included 6-10 grains of active dry yeast (Red star C2751) in each vial 24 hours before the experiment. The next day, flies were briefly anesthetized using CO_2_, and while immobilized, were manually transferred to every oviposition well using a reagent scoopula (Fisherbrand^TM^ Scoopula^TM^ Spatula C14357Q) and small paint brush. Each female was paired with 2 males in every oviposition chamber. We ran the assay for 24 hours in an incubator chamber under the same conditions as rearing.

We assayed 155 RILs from panel B of the DSPR for oviposition preference. Each RIL was assayed across 4 independent replicate chambers, with each chamber stocked with 15 females (paired with 30 males) for a target of 60 females per RIL across four replicates. Assays were conducted in batches, with multiple RILs tested in parallel. To avoid systematic bias due to batch or day effects, RIL identity was randomly assigned to plates to minimize within batch positional and tray effects, and the side assigned to Cu versus control substrate was randomized across batches to control for within chamber positional bias. Replicate counts (3 to 4) varied slightly due to differences in the availability of mated females from some DSPR stocks. After the oviposition period, adults were removed, and dead females were excluded from egg counts. Female mortality during the 24-hour assay was low overall (47 dead out of 9,090 females assayed; 0.52%), affecting 34 of 155 RILs (mean 0.30 deaths per RIL, range 0-7) and 40 of 606 replicate chambers (mean 0.08 deaths per chamber, range 0-6). Eggs were manually counted from photos taken using a high magnifying DSLR camera (Nikon D780, SIGMA 105mm 1:2.8 DG MACRO HSM), carefully distinguishing eggs on the Cu media versus on the control media using the Cell Counter plugin developed for ImageJ software (Schneider et al. 2012).

To quantify oviposition preference in response to Cu, we calculated oviposition preference for each replicate vial using the formula:

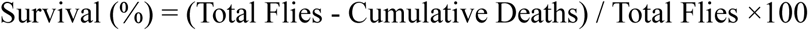

As for feeding preference, negative values indicate avoidance of Cu and positive values indicate preference for Cu. We excluded replicate chambers with fewer than five total eggs across both substrates and RILs retaining fewer than three valid replicates, resulting in a final analyzed dataset of 149 RILs.

To assess whether oviposition preference differed among genotypes, we performed a one way ANOVA with RIL as a fixed effect and oviposition preference index (PI) as the response variable (PI ∼ RIL). Prior to analysis, model assumptions were evaluated using residual Q-Q plots and histograms. Residuals were approximately normal, with no major departures requiring transformation, so PI was analyzed on its original scale.

We estimated the broad sense heritability (H^2^) of oviposition preference as the proportion of total phenotypic variance explained by among-RIL differences. Variance components were estimated using a linear mixed model with RIL as a random effect (PI ∼ 1 + (1 | RIL) using the *nlme* package (Pinheiro et al. 2025). Because the oviposition assay was conducted across 19 assay days, with approximately 9-10 RILs tested per day, most RILs were measured within a single batch rather than across multiple batches. Therefore, RIL identity and assay day were partially confounded. To evaluate whether this affected the heritability estimate, we also fit a replicate level mixed model with both RIL and assay day as random effects (PI ∼ 1 + (1 | RIL) + (1 | day)), using restricted maximum likelihood (*lme4* package (Bates et al. 2015)). From this model, we calculated a conservative H^2^ estimate by treating assay day variance as part of the non-genetic variance, and a second estimate by partitioning assay day variance out of the denominator.

To test whether variation in oviposition preference was associated with founder haplotype variation in the DSPR, we performed QTL mapping using the DSPRqtl (version 2.0-5) and DSPRqtlDataB (version 2.0-1) packages (King et al. 2012). For QTL mapping, per RIL mean preference was √(PI + 1) transformed to stabilize variance across the founder haplotype model and to maintain consistency with the feeding QTL analysis, which used the same transformation. In contrast, the RIL ANOVA described above was performed on untransformed PI because residual diagnostics indicated that model assumptions were adequately met. All analyses were performed in R (R Core Team 2025).

### Physiological response to copper

Adult males and females (3-5 days post-eclosion) were sorted under light CO_2_ anesthesia in groups of 15 flies and allowed to recover for 24 h. We set up 3 independent replicate vials for every combination of RILs, sex (male, female), control treatment (no copper) and Cu treatment (2, 5, & 10 mM CuSO_4_). Because the full panel could not be assayed in a single run, the experiment was conducted in two batches of 10 RILs each, for a total of 20 RILs. All sex and treatment combinations were balanced within each batch. After recovery, flies were transferred (without anesthesia) to semidefined medium recipe (Backhaus et al. 1984) vials supplemented with the designated concentration of CuSO_4_ (control, 2, 5, & 10 mM). Each vial was assigned a unique barcode and tracked through the assay. We recorded the initial number of flies per vial on Day 0 to confirm that each vial began with 15 individuals and to record any flies that had died during handling (excluded from data analysis). We recorded mortality every 24 hours until LT_50_ was reached in all Cu treated vials. At that point, the experiment was concluded, even if flies were still alive in control (0 mM) vials, as our analysis focused exclusively on Cu exposed treatments. To ensure that treatment conditions were not impacted by larval activity or bacterial contamination, the food medium was replaced with freshly prepared same food vials 3 times a week.

To quantify physiological resilience in RILs under Cu stress, we estimated the median lethal time (LT_50_) for each vial using time series survivorship data. Daily mortality rates were compiled from vials that were scanned daily using a barcode data logger (AML striker). Raw mortality data were adjusted to account for individuals that died from initial handling, and survival percentages were calculated daily as:

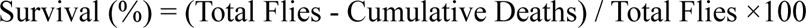

LT_50_ values were determined in R using a custom function that applied linear interpolation for each vial as the time point at which survival crossed 50%, using the two nearest surrounding survival values (Robertson and Preisler 1992). The interpolation function and downstream mixed effects models are provided in Supplementary File S1.

To test the effect of Cu treatment on LT_50_ while accounting for genetic background, we fit a linear mixed effects model LT_50_ ∼ Treatment × Sex + Batch + (1 | RIL); using the *lme* function from the *nlme* package (Pinheiro et al. 2025). RIL was treated as a random effect to capture inter RIL variability, and batch was included as a fixed effect to account for differences between assay runs. Because residual variance decreased with Cu dose, based on Levene’s test of residuals from an initial homoscedastic model (p < 0.001), we modeled unequal residual variance across treatment levels using a *varIdent* variance structure (Pinheiro and Bates 2000). This improved model fit relative to the homoscedastic model (ΔAIC = 19). Model residuals were evaluated using Q-Q plots and the Shapiro-Wilk test (Fox 2015). Fixed effects were tested using Type II F tests with Satterthwaite degrees of freedom from the *nlme* model and were cross checked with Kenward-Roger F tests from the homoscedastic model using the *car* package (Fox and Weisberg 2019). Post-hoc pairwise comparisons among treatments were performed using EMMs with Tukey adjusted p-values (*emmeans* package (Lenth 2023)). Significant groupings were visualized with compact letter displays (CLD) using *multcompView* (Hothorn et al. 2008). The intraclass correlation coefficient obtained from the fitted model was used to estimate the proportion of LT_50_ variability associated with RIL. All analyses were performed in R (R Core Team 2025).

### Phenotypic correlations among copper response components

To test whether the three components of Cu response (feeding avoidance, oviposition avoidance, and physiological tolerance) were correlated among RILs, we performed pairwise correlation analyses across DSPR RILs. These included female feeding preference versus oviposition preference, and each behavioral trait versus LT_50_ at three CuSO_4_ doses. We used only female feeding preference for these correlations because oviposition is a female specific trait.

For each comparison, we used RIL means as the unit of analysis (one data point per RIL), with values averaged across replicate measurements within each trait. Only RILs measured for both traits in a given pairwise comparison were retained. The feeding by oviposition comparison included 143 RILs, whereas behavior by LT_50_ comparisons used the shared subset of 18 RILs. Because feeding preference was assayed at 2 mM CuSO_4_ and oviposition preference at 10 mM CuSO_4_, each RIL’s preference index was treated as a summary estimate of Cu avoidance behavior within that assay context rather than as a dose matched measurement. For each pairwise comparison, we fit a linear regression with lm() and calculated Pearson’s r with 95% confidence intervals using cor.test() in R (R Core Team 2025). Spearman correlations and a tercile based concordance analysis, including Fisher’s exact test and Cohen’s kappa calculated with the *psych* package (Revelle 2026), are reported in Supplementary Table S2. Because the physiology panel was substantially smaller than the behavioral panels, correlations involving LT_50_ are reported with 95% confidence intervals and should be interpreted cautiously given the limited statistical power.

## Results

### Feeding aversion to Cu varied among DSPR RILs

Aversion to Cu contaminated food was variable among the DSPR RILs (DSPR RIL: F _(196,10101)_ = 14.33, P < 0.0001; Figure 1A, B) and heritable in both sexes (H^2^ = 88%). The majority of RILs displayed strong avoidance of Cu contaminated food (Figure 1), with females avoiding Cu more strongly than males (Sex: F _(1,10101)_ = 349.70, P < 0.0001; Figure 1A and B). We detected a minor interaction between DSPR RIL and Sex (F _(181,10101)_ = 2.25, P < 0.0001; Figure 1C), but overall, male and female feeding preference was significantly correlated (F _(1,180)_ = 223.7; P < 0.0001, adjusted R^2^ = 55.2%, r = 0.74, 95% CI 0.67–0.80, P < 0.001; Figure 1D).

**Figure 1.**
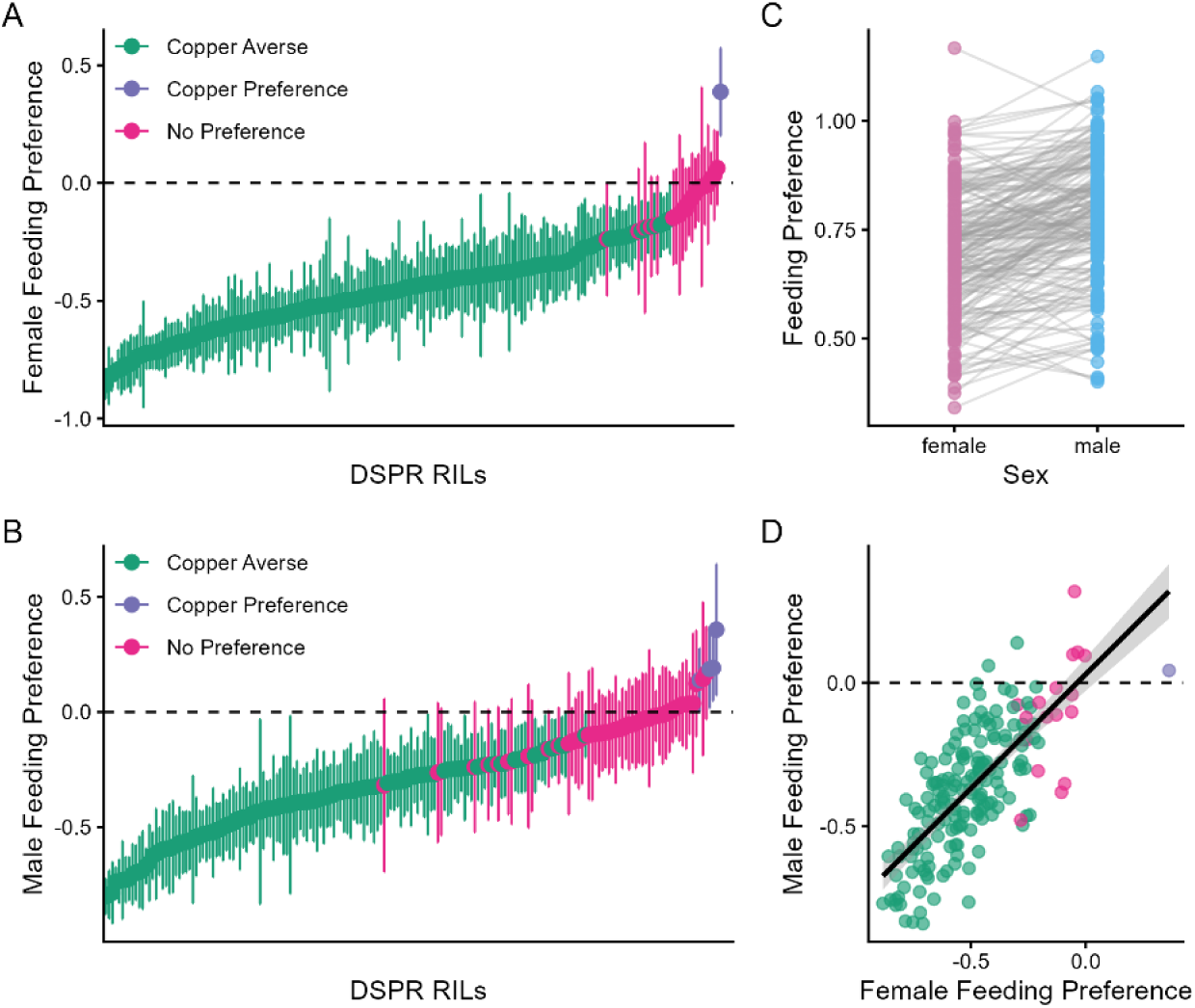
DSPR RILs show heritable variation in Cu feeding avoidance, and females avoid Cu more strongly than males. (A) Female feeding preference. Females tended to avoid Cu (green), with a small subset of RILs consuming both control and Cu food at a similar level (pink). One RIL preferred Cu food (purple). (B) Male feeding preference. Males avoided Cu less strongly than females, with more RILs choosing control or Cu food at an indistinguishable rate (pink). Four DSPR RILs preferred Cu food (purple). In 1A and1 B, points show mean DSPR response ± 95% CI. (C) Female-male comparison of RIL mean feeding preference; lines connect the same RIL. A minor but significant Sex × RIL interaction was detected. (D) RIL level correlation between male and female feeding preference (Pearson r = 0.74). Values in Fig 1 A, 1B, and 1D are shown on the raw preference scale and in 1C are on the transformed scale. Dashed lines (A, B, D) indicate no preference. DSPR RILs are ordered independently by mean response in 1A and 1B.

We performed QTL mapping to test whether variation in Cu feeding preference was associated with founder haplotype differences in the DSPR. No QTL were associated with variation in female Cu feeding preference, but we detected one QTL that explained 17.7% of the variation in male Cu feeding preference (Figure 2A). The structure of the DSPR mapping panel allows the effects of the founder alleles to be estimated using the DSPRqtl package. Estimated effects of each founder haplotype with sufficient representation (> 5 RILs) suggest the presence of two distinct allele classes (Figure 2B). Other factors, including unique alleles with similar effects, non-additive founder haplotype effects, and variation in founder haplotype representation, may influence this estimates, particularly given our small sample size.

**Figure 2.**
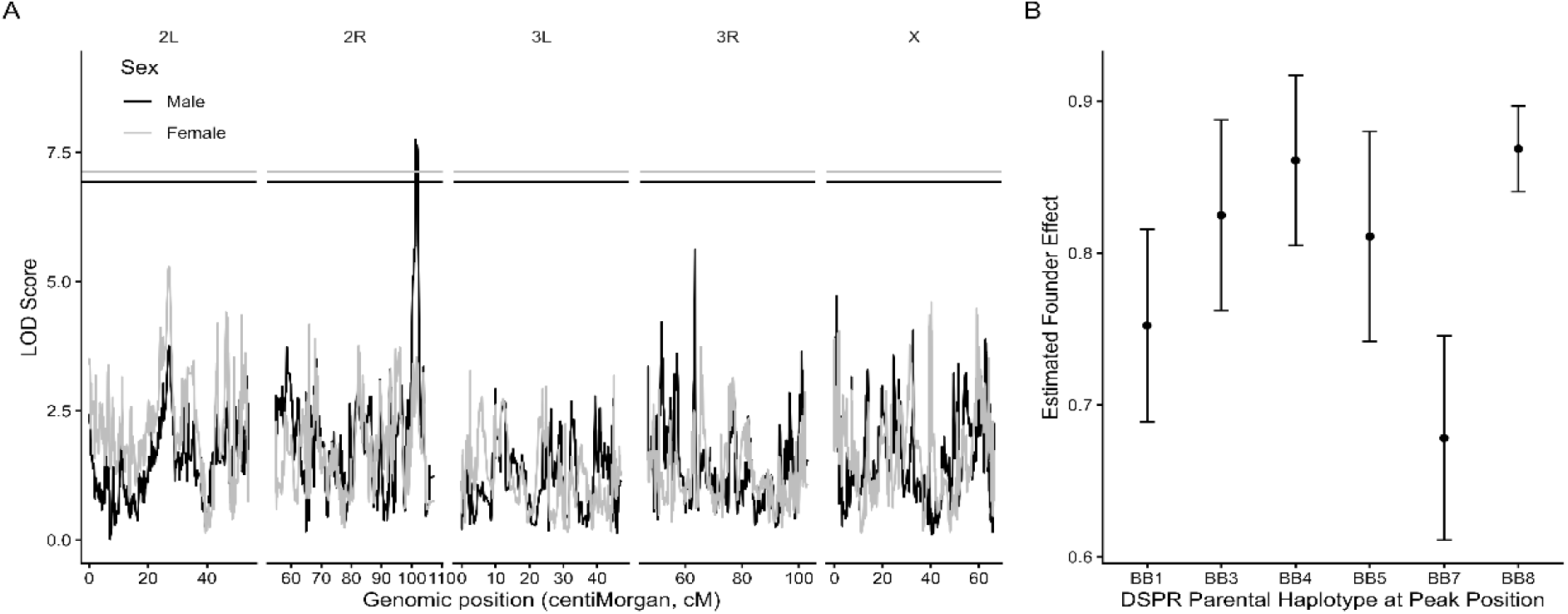
A male-specific QTL on chromosome 2R contributes to variation in Cu feeding preference. (A) Genome-wide LOD profiles for male (black) and female (grey) feeding preference across the five major chromosome arms (2L, 2R, 3L, 3R, X), plotted against genomic position in centimorgans (cM). Horizontal lines indicate permutation derived significance thresholds (α = 0.05, 1,000 permutations). (B) Estimated founder haplotype means at the 2R peak for male feeding preference. Points show estimated effects ± 95% CI on the √(feeding preference + 1) transformed scale. A value of 1 corresponds to no feeding preference; values below 1 indicate Cu avoidance, with lower values representing stronger avoidance. BB2 and BB6 were excluded because they were represented by fewer than five.

Using a 2 LOD drop, the male feeding QTL peak (lower bound: 2R:18780000bp; upper bound: 2R:19060000) encompassed a genomic region with 39 genes, 34 of which are protein coding (Table S1). Of these genes, promising candidates include *sano* (FBgn0034408) and *Jheh1*, *Jheh2*, and *Jheh3* (FBgn0010053, FBgn0034405, and FBgn0034406). The gene *sano* has been linked to olfactory development and decision making in a two choice aversive conditioning experiment (Walkinshaw et al. 2015). The three juvenile hormone epoxide hydrolases have been recently linked to regulation of the gene expression response to lead (Wang et al. 2025).

### Oviposition avoidance varied among RILs but yielded no detectable QTL

We quantified oviposition preference for Cu containing versus control substrate across DSPR recombinant inbred lines (RILs) using a preference index (PI) and total egg counts. Across the full dataset, fecundity was highly variable (total eggs range: 9-943 per replicate). After filtering to retain lines with adequate replication (≥3 replicates per RIL), the final oviposition dataset included 149 RILs.

Oviposition preference differed significantly among RILs, indicating substantial genetic variation in reproductive substrate choice under Cu exposure (ANOVA: F _(148, 446)_ = 11.88, p < 0.001) (Figure 3). Mean PI values per RIL spanned a broad range, with many genotypes showing strong Cu avoidance and a subset exhibiting relatively weak avoidance or preference near or above zero. Finally, variance partitioning using a random effects model on RIL means indicated high broad sense heritability of oviposition preference across RIL averages (H^2^ ∼ 0.88), consistent with strong among line (genetic) contributions relative to within line variation in this assay. Because most RILs were assayed within a single batch (9-10 RILs per batch across 19 assay days), we refitted the model on replicate level data with assay day as an additional random effect to confirm this estimate was not driven by batch structure. Assay day accounted for only 8% of phenotypic variance, compared with 65% explained by RIL, and heritability remained substantial after accounting for batch structure (H^2^ = 0.65 when day variance was treated as non-genetic variance; H^2^ = 0.71 when day variance was partitioned out). Therefore, the high heritability of oviposition preference is unlikely to be an artifact of assay day effects.

**Figure 3:**
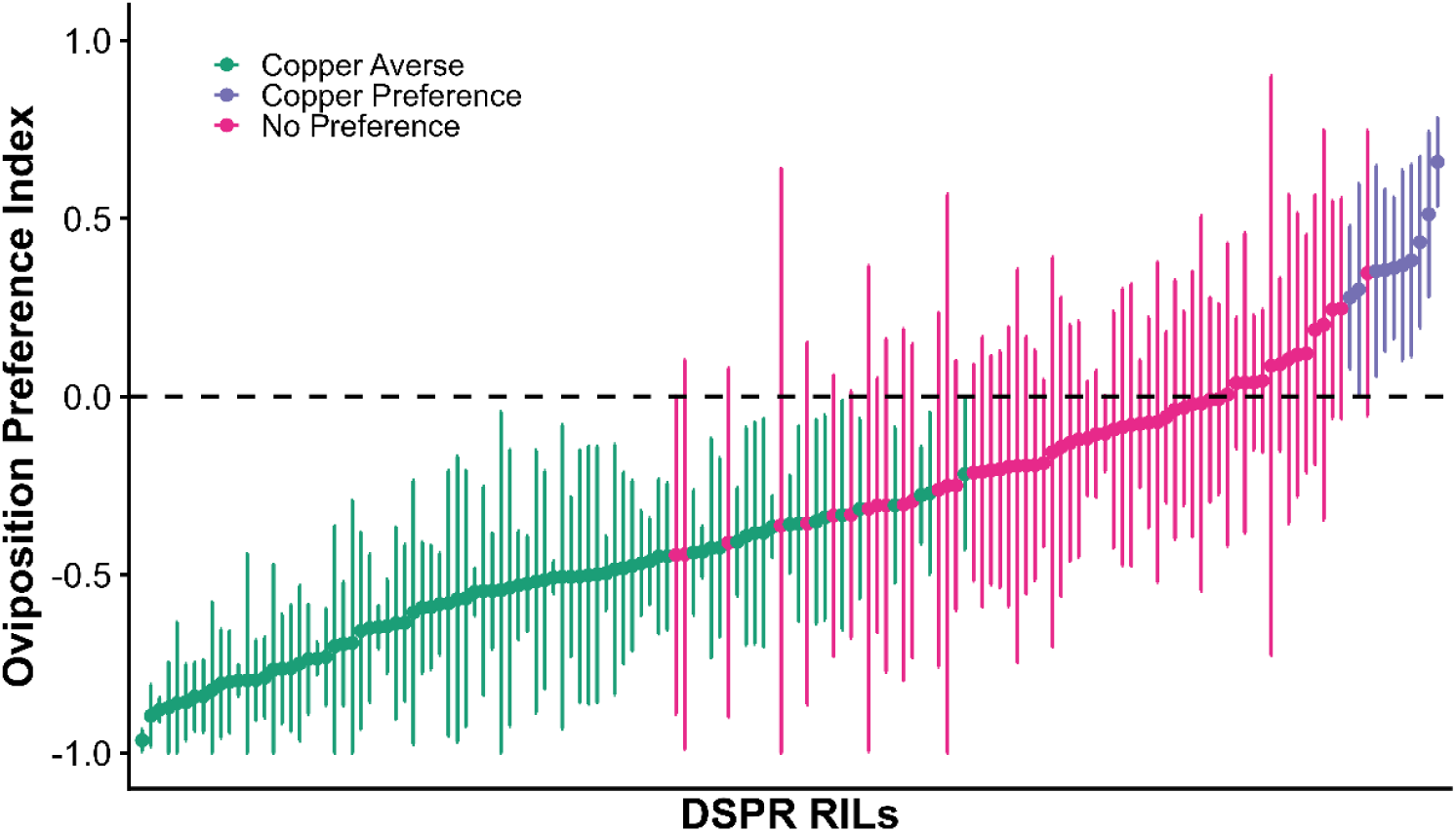
DSPR RILs vary in oviposition response to Cu, with most avoiding Cu substrate. Oviposition preference index for 10 mM CuSO_4_ vs. control substrate across DSPR RILs. Points show RIL mean ± 95% CI. RILs are colored by group based on whether the 95% CI excluded zero; Cu averse (green), no preference (pink), or Cu preferring (purple) and ordered along the x-axis by mean preference. The dashed line at PI = 0 denotes no preference.

To test whether heritable variation in oviposition preference could be attributed to founder haplotypes, we performed QTL mapping using the √(PI + 1) transformed mean preference for each RIL as the response variable in the DSPRqtl framework. No QTL exceeded the genome wide significance threshold on any chromosome arm in either the female only or combined analyses (Figure S3). Across the genome, peak LOD scores for oviposition preference remained below the permutation derived threshold, and no peaks overlapped with the male feeding QTL on chromosome arm 2R. Although no significant QTL were detected, oviposition preference exhibited high broad sense heritability (H² ∼ 0.88). Given the panel size used here (N= 149 RILs), this pattern is consistent with a polygenic architecture in which variation is distributed across multiple loci of small effect rather than concentrated at a few large effect loci. In contrast, male feeding preference showed a detectable QTL on 2R explaining 17.7% of the phenotypic variance, suggesting differences in the genetic architecture underlying these traits.

### Copper reduces survival in a dose dependent, RIL variable manner

Cu exposure significantly reduced adult survival time in a dose dependent manner (Figure 4). At the lowest tested dose, mean LT_50_ across RILs was 31.2 days at 2mM CuSO_4_. LT_50_ declined to 26.9 days at 5 mM and to 20.1 days at 10 mM CuSO_4_, with all pairwise Tukey contrasts significant (p < 0.0001). This dose dependent reduction in survival was consistent across both sexes, although the Treatment × Sex interaction was at the margin of significance (F _(2,335)_ = 3.10, p = 0.046). Females and males had similar LT_50_ values at 2 mM (30.4 vs. 32.1 days) and 5 mM CuSO_4_ (26.7 vs. 27.0 days), but females survived slightly longer than males at 10 mM CuSO_4_ (20.7 vs. 19.5 days). LT_50_ also varied substantially among RILs at each Cu dose (Figure 4). The among RIL variance component (σ² RIL = 19.9) accounted for 49.5% of the residual variance in LT_50_, indicating that genetic background contributed meaningfully to physiological tolerance. At 10 mM CuSO_4_, the most tolerant RIL survived approximately 2.3 times longer than the least tolerant RIL, based on LT_50_ estimates of 25.7 and 11.4 days, respectively. Because the physiology panel was limited to 20 RILs, we did not perform QTL mapping for LT_50_ in this dataset.

**Figure 4.**
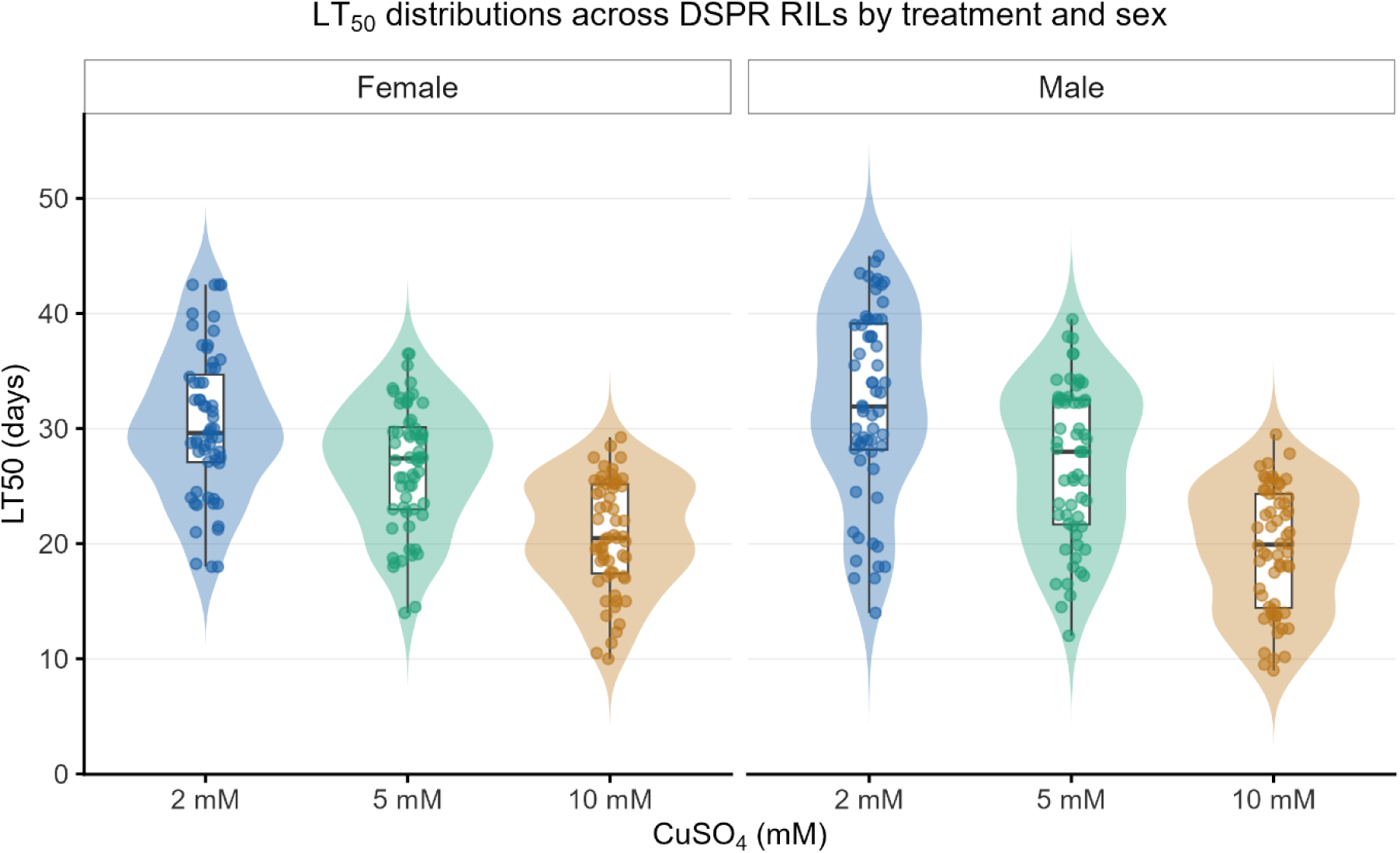
Copper reduces LT_50_ in a dose dependent, sex, and RIL dependent manner. Distribution of vial level LT_50_ (median lethal time, days) across 20 DSPR RILs at three Cu doses (2, 5, and 10 mM CuSO_4_), separated by sex (n = 60 vials per treatment × sex cell; 3 replicate vials per RIL × sex × treatment combination). Violins show the distribution across vials, boxplots the median and interquartile range, and points individual vials. LT_50_ declined with dose in both sexes (F _(2,335)_ = 201.5, p < 0.0001), with a significant Treatment × Sex interaction (F _(2,335)_ = 3.10, p = 0.046).

### Feeding, oviposition, and physiological tolerance show weak correlations across RILs

Across the 143 RILs assayed for both feeding and oviposition, female feeding preference and oviposition preference index (PI) were not significantly correlated (R^2^ = 1.37%, F _(1,141)_ = 1.96, p = 0.163; Figure 5A). Rank based tercile classification showed a similar lack of association: 47 of 143 RILs (32.9%) fell into the same Cu sensitivity tercile for both traits, close to the 33.3% expected under independence (Fisher’s exact test, p = 0.997; Cohen’s kappa = −0.01, 95% CI [−0.12, 0.11]; Table S2). A further 30 RILs (21.0%) were classified in opposite terciles, indicating that they were relatively Cu sensitive in one behavioral assay but relatively Cu tolerant in the other.

**Figure 5.**
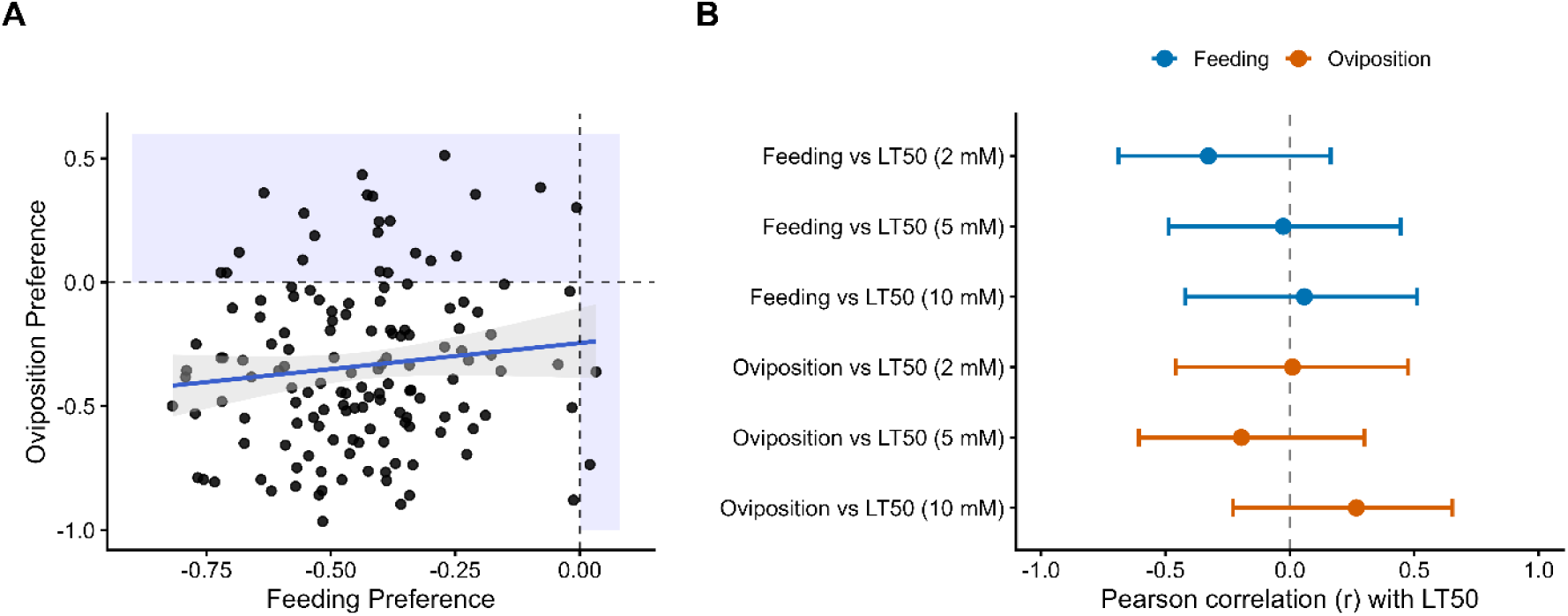
Feeding, oviposition, and physiological tolerance are largely independent among DSPR RILs. (A) Female feeding preference and oviposition preference index (PI) are not significantly correlated across the 143 RILs assayed for both traits (R2 = 1.37%, F _(1,141)_ = 1.96, p = 0.163). Points are RIL means; solid line is the OLS regression with 95% confidence band. Dashed reference lines mark neutral preference on each axis; shaded quadrants indicate the Cu preference region. (B) Pairwise Pearson correlations between behavioral traits (female feeding preference, oviposition PI) and physiological tolerance (LT_50_) at three Cu doses, across the 18 RILs measured for all three phenotypes. Points show point estimates; horizontal bars are 95% CI; dashed reference line at r = 0. All confidence intervals overlap to zero, consistent with independent among RIL variation in the behavioral and physiological components of the Cu response.

Across the 18 RILs measured for all three phenotypes, neither female feeding preference nor oviposition preference were significantly correlated with LT_50_ at any Cu dose, and all six 95% confidence intervals included zero (Figure 5B; Table S3). Feeding preference showed a weak, non-significant negative relationship with LT_50_ at 2 mM CuSO_4_ (r = −0.33, 95% CI [−0.69, 0.17], p = 0.19), but this relationship was not observed at higher doses (5 mM: r = −0.03; 10 mM: r = 0.06). Oviposition preference was similarly uncorrelated with LT_50_ across doses (|r| ≤ 0.27, all p ≥ 0.28).

## Discussion

We set out to investigate whether Cu resistance in *D. melanogaster* arises from behavioral avoidance, physiological tolerance, or both, and whether these components share a common genetic basis. Our findings demonstrate that feeding avoidance, oviposition avoidance, and physiological tolerance are heritable traits but vary independently across RILs, with no common QTL identified among traits. Broad sense heritability was high for both feeding preference (H^2^ ∼ 0.88) and oviposition preference (H^2^ ∼ 0.88), indicating that genetic variations among RILs explain the phenotypic variance in each trait. Nevertheless, the two traits were only weakly correlated across RILs (R^2^ = 1.37% and only 32.9% of RILs fell into the same Cu sensitivity tercile for both traits no different from independence), indicating that the genetic factors that affect feeding avoidance are likely different from those influencing oviposition site choice. Together, these results suggest that different components of Cu resistance can evolve independently.

A small number of DSPR RILs consumed Cu-supplemented food at rates similar to or higher than control food. This pattern is consistent with previous characterizations of Cu resistance and behavioral responses to Cu as complex and genotype-dependent (Bahadorani and Hilliker 2009; Everman et al. 2021). We interpret these RILs as exhibiting a failure of avoidance rather than true preference, consistent with observations in Cu sensitive genotypes (Everman et al. 2021). We did not directly evaluate the sensory or motivational basis of this pattern, so this interpretation remains hypothetical. Targeted follow up work comparing gustatory receptor expression or feeding circuit activity between Cu-non-avoidant and strongly Cu-avoidant RILs would help distinguish a true sensory deficit from alternatives such as elevated metabolic demand or shifted satiety thresholds. Both feeding output and detoxification investment are known to be modulated by internal physiological state in *Drosophila* and other insects (Castañeda et al. 2009; Wang and Wang 2019), but whether such differences underlie variation in Cu avoidance remains untested.

Regardless of mechanism, the organismal consequence is the same. Failing to avoid Cu means ingesting more Cu. However, feeding preference was not significantly associated with LT_50_ in our RIL panel, suggesting that behavioral avoidance and physiological tolerance may contribute independently to Cu response. Increased Cu ingestion elevates internal metal burden, disrupting redox homeostasis through reactive oxygen species production and oxidative damage to cellular macromolecules (Stohs and Bagchi 1995). Together, these effects suggest that genotypic variation in feeding avoidance may have fitness consequences beyond the behavioral endpoint itself.

Notably, we detected a single QTL on chromosome arm 2R explaining 17.7% of the variance in male feeding preference, with no significant QTL detected in females or for oviposition preference. The 2 LOD support interval spans 39 genes, 34 of which are protein coding (Table S1). Among these, the juvenile hormone epoxide hydrolase genes *Jheh1*, *Jheh2*, and *Jheh3* are particularly interesting candidates. These 3 *Jheh* paralogs are clustered together on chromosome 2, where they function in juvenile hormone metabolism, xenobiotic detoxification, and stress response (Rogalski et al. 2025). At this locus, *Jheh1* and *Jheh2* are induced under xenobiotic and oxidative stress (Deng and Kerppola 2014), and lead (Pb) exposure also induces both genes in the *D. melanogaster* midgut through an *MTF1*/*Nrf2*/*JNK* mediated heavy metal response (Wang et al. 2025). Another candidate in the interval is *sano/serrano* (CG12758). This gene was previously identified in a panneuronal RNAi screen in which knockdown reduced olfactory memory performance in *Drosophila* (Walkinshaw et al. 2015). This makes *sano* potentially relevant to sensory guided feeding decisions, although its primary annotated role is in planar cell polarity. Because our panel size (n = 197 RILs) provides sufficient power to detect a QTL of this effect size, it does not allow fine mapping to individual genes. Functional validation such as RNAi knockdown or allele replacement experiments targeting the *Jheh* cluster or the *sano* will be required to directly test their roles in Cu related feeding behavior.

While female *Drosophila* generally avoid ovipositing on heavy metal contaminated substrates to protect offspring (Bahadorani and Hilliker 2009; Gou et al. 2014), oviposition site choice is an active decision making behavior with females evaluating substrate cues for each egg they lay (Yang et al. 2008). The small subset of RILs in our panel that laid eggs on Cu substrate at rates comparable to control is therefore biologically notable. Paralleling our interpretation of the feeding outliers, we suggest these RILs reflect a failure of avoidance rather than true Cu preference. Cu exposure has been shown to impair locomotor activity, damage dopaminergic neurons in *Drosophila* (Bonilla-Ramirez et al. 2011), and to reduce fecundity at sublethal doses (Budiyanti et al. 2022), suggesting multiple physiological pathways through which Cu could disrupt the sensory system underlying substrate evaluation. Therefore, neural circuits involved in this decision are plausible targets. However, identifying which circuits are affected in our outlier strains would require direct neural or transcriptomic measurements that we have not yet performed. The contrast between the feeding and lack of oviposition QTL results suggests that these behaviors are governed by independent genetic architectures. We detected a male-specific feeding QTL on chromosome arm 2R, but none in females and none for oviposition preference, despite similarly high heritability for both traits (H^2^ ∼ 0.88). Such sex dependent genetic effects on metal response were also recently reported for nickel and Cu induced mortality in *D. melanogaster*, where line, sex, and their interaction all shaped survival (Hutchings et al. 2026). Our QTL results suggest that feeding avoidance, at least in males, may involve loci of moderate to large effect, whereas oviposition avoidance may be influenced by many loci of smaller effect. Combined with the near zero phenotypic correlation between feeding and oviposition preferences (R^2^ = 1.37%), the absence of detectable oviposition QTL is consistent with these behaviors being genetically distinct, though we cannot exclude the possibility that greater statistical power would reveal some shared loci.

The phenotypic independence of feeding and oviposition preferences (*R*^2^ = 1.37%) indicates that decisions governing ingestion of toxins versus deposition of eggs on toxins operate through genetically independent pathways. This is consistent with prior work showing that Cu resistance in *D. melanogaster* integrates behavioral and physiological components, with feeding behavior negatively correlated with adult Cu resistance across DSPR strains (Everman et al., 2021). Similar decoupling has been reported in other systems. For example in *D. melanogaster*, behavioral avoidance and physiological resistance to malathion were genetically uncorrelated across wild strains (Pluthero and Threlkeld 1981) and in the diamondback moth (*Plutella xylostella*), selection for larval physiological resistance to *Bacillus thuringiensis* toxins did not alter adult oviposition preference (Groeters et al. 1992).

Another possible explanation for this decoupling is tissue specific regulation. Cu responsive gene expression differs sharply between head and gut tissue, and the metallothionein gene *MtnA* is regulated by a *cis*-eQTL in gut but a *trans*-eQTL in head (Everman and Macdonald, 2024). Strains may therefore harbor regulatory variants that preserve gut mediated avoidance during feeding while independently compromising processing of oviposition cues, or vice versa. The independent genetic control of these pathways allows them to respond to selection separately, giving populations modular behavioral flexibility in heterogeneous toxic environments.

### Linking behavior, physiology, and the modular architecture of Cu resistance

Our physiological assays showed a clear dose dependent reduction in LT_50_ with increasing Cu concentration, with substantial variation among DSPR RILs (Figure 4), with RIL identity accounting for 49.5% of the modeled variance. This suggests that genetic background contributes to physiological tolerance. We selected behavioral assay concentrations to maintain >90% adult survival, allowing us to measure behavioral responses rather than mortality. As a result, survival is a relatively coarse readout at these doses. Behavioral effects can precede mortality, and sublethal Cu exposure is known to impair learning and locomotion in *Drosophila* before survival is affected (Tran Thanh et al. 2024). RILs that fail to avoid Cu through feeding or oviposition are therefore likely to experience fitness costs that are not captured by LT_50_ alone, including potential downstream effects on offspring condition (Mu et al. 2021) and larval immune function (Nanda et al. 2019).

This brings us back to the question we set out to address. If behavioral avoidance and physiological tolerance shared a common genetic basis, we would expect RILs exhibiting strong Cu avoidance to also show high physiological tolerance and overlapping QTL for these traits. Instead, feeding and oviposition preferences were highly heritable, while LT_50_ showed substantial genetic variation among RILs, yet these traits were uncorrelated. Across the 18 RILs measured for all three phenotypes, there were no significant correlations between feeding preference or oviposition preference with LT_50_ at any Cu doses. This finding suggests that such responses are at least somewhat independent, although the small sample size of RILs limits our ability to detect weaker correlations. The QTL results provide additional support. The only QTL detected in our study, the male feeding QTL on chromosome arm 2R (18.81 Mb), also did not overlap with the physiological Cu resistance QTL previously mapped in the same DSPR B panel by Everman et al. (2021). Their two 2R resistance QTL map proximally (1.4-6.6 Mb), approximately 38 cM from our distal feeding QTL. This non-overlap provides further evidence that feeding avoidance and physiological tolerance are influenced by distinct loci. Together, these results are consistent with a model in which Cu resistance in *D. melanogaster* is composed of multiple, partially independent components, including the ability to detect and avoid Cu during feeding, evaluate substrates during oviposition, and physiologically tolerate internal Cu exposure. This structure suggests that selection can act on these components independently, allowing populations to evolve different combinations of behavioral and physiological responses depending on the selective environment.

### Limitations and next steps

Several caveats are worth stating directly. First, the physiology panel was small (20 RILs), so we did not estimate quantitative heritability for tolerance or map QTL for it in this dataset; instead, we compared our male feeding QTL against the physiological Cu resistance QTL mapped in the same DSPR B panel strains by Everman et al. (2021), which showed no overlap on 2R. A larger panel would still be needed to determine whether physiological tolerance has its own QTL architecture beyond the resistance loci already mapped in these strains. Second, the *Jheh1-3* cluster and *sano* within our 2R interval is a plausible candidates based on prior work in metal response gene regulation (Rogalski et al. 2025; Wang et al. 2025) and olfactory memory function (Walkinshaw et al. 2015), respectively, but it has not been directly validated in a Cu context, and the interval contains 34 protein-coding genes. Third, we did not detect significant QTL for oviposition preference despite high heritability (H² ∼ 0.88), which we interpret as evidence for a highly polygenic architecture rather than as a null result; this should be tested in a larger panel. Finally, the mechanistic hypotheses we propose, including sensory deficits underlying failure of avoidance and tissue specific regulatory variation contributing to the uncoupling of feeding and oviposition, require direct experimental tests that we have not yet performed.

## Conclusion

Behavioral and physiological responses to Cu in *Drosophila melanogaster* are heritable but likely genetically modular. Feeding avoidance, oviposition avoidance, and physiological tolerance are three semi-independent components of Cu resistance, with no detectable phenotypic correlation across DSPR RILs. The QTL we mapped (male feeding, chromosome arm 2R, 17.7% of variance) provides a starting point for identifying causal variants underlying behavioral Cu avoidance, and is genetically distinct from the physiological Cu resistance QTL previously mapped in the DSPR (Everman et al. 2021). Our results suggest that selection may act on different axes of Cu resistance independently, with implications for how natural populations adapt to heavy metal contamination, where fitness depends on whether organisms can detect, avoid, or physiologically tolerate the stressor.

## Author Contributions

**Md Meftahul Zannat**: Conceptualization, Methodology, Investigation, Formal analysis, Data curation, Visualization, Writing: Original draft, review & editing. **Jordan C. Jones**: Investigation, Data curation, Writing: Components of original draft, review, & editing. **Maggie Ridgway**: Methodology, Writing: review & editing. **Elizabeth R. Everman**: Conceptualization, Resources, Supervision, Funding acquisition, Project administration, Formal analysis, Writing: Review & editing.

## Data availability

All data and relevant R code used in this study will be available from Dryad (DOI: https://doi.org/10.5061/dryad.83bk3jb8s). R scripts for the feeding and oviposition analyses, the physiology LT_50_ pipeline, and phenotypic correlations are provided in **Supplementary File S1**.

## Acknowledgements

We thank Kazzrie Arnold for expert assistance with fly sorting and are grateful to Dr. Ingo Schlupp and Dr. Dave Hambright, Claire Grimmett, Chuck Miller, for reviewing this manuscript. NIH NIEHS 4R00ES033257-03 funded this work.

